# Unbiased proteomics following inflammasome activation identifies caspase targets in primary intestinal epithelial cells

**DOI:** 10.64898/2026.04.20.719683

**Authors:** Alexis R. Gibson, Ivo Diaz Ludovico, Geremy C. Clair, Chelsea M. Hutchinson-Bunch, Joshua Adkins, Isabella Rauch

## Abstract

Inflammasomes are cytosolic innate immune sensors that, once activated by a pathogenic threat, lead to activation of the inflammatory Caspase-1. Inflammasome activation and its consequences have been studied extensively in myeloid cells and in overexpression systems. Recent studies have identified cell type specific effects that are not fully explained by the known cleavage targets of Caspase-1. Here, we identified targets of caspase cleavage using mass spectrometry in primary intestinal epithelial cells by specifically activating the NAIP-NLRC4 inflammasome. We have taken an unbiased approach and developed a novel method for analyzing mass spectrometry data for evidence of caspase activity. Our approach can also be applied to existing proteomic datasets to establish the presence of caspase activity under various biological conditions. These results lay the groundwork for future studies on mechanisms of caspase-induced processes such as intestinal epithelial cell extrusion.

## Introduction

Caspases (CASP) are a family of proteases that act as central executioners of inflammation and apoptosis, implicated in the pathogenesis of immune, cardiovascular, and metabolic diseases, including cancer (1-5). The inflammatory CASP-dependent action is regulated by inflammasomes, multiprotein complexes formed after the recognition of intracellular infection or danger signals in the cytosol of a cell. Inflammasome activation canonically leads to activation of the inflammatory programmed cell death executor protease CASP1, cleaving and activating cytokines and the pore-forming protein Gasdermin D (GSDMD), leading to a programmed lytic cell death called pyroptosis.

Previous studies have sought to identify inflammatory caspase substrates in immortalized cell lines, exclusively those derived from hematopoietic cells (6-8). Yet inflammasome activation leads to striking cell type specific responses, such as rapid cell extrusion in intestinal epithelial cells (IEC) (9). Cell extrusion involves epithelial cell contraction (10) as well as coordinated cytoskeleton rearrangement (11, 12), but the exact mechanism of cell extrusion after pyroptosis is unclear. Similarly, caspase substrate proteins are not yet fully characterized, potentially obscuring their roles in targeted therapeutic strategies for inflammatory processes. However, there are some well-characterized examples of caspase substrates that could play more important roles in certain cell types. For instance, CASP7, a known CASP1 substrate (7), was recently shown to execute the late stages of extrusion of IEC (13). Stromal cells, such as IEC, also have dramatically different gene expression profiles from immune cells. Therefore, it is likely that there are substrates of caspases that are unique to IEC and contribute to efficient extrusion, with many yet to be discovered.

Currently, powerful methods are available for the detection of caspase substrates (14). However, these techniques often require time-consuming procedures, specialized equipment, and costly reagents. Moreover, despite their utility, identified targets still require additional validation steps. For example, several approaches rely on N-terminomics, which involves enrichment or labeling steps followed by deep mass spectrometry analysis, yet findings obtained by these methods often still need further confirmation.

We hypothesized that a modern data-dependent analysis (DDA) label-free quantification (LFQ) mass spectrometry pipeline could be used to identify caspase targets at the peptide level. To test this idea, we took advantage of our system to specifically activate the neuronal apoptosis inhibitory protein (NAIP)/NLR family caspase activation and recruitment domain-containing protein 4 (NLRC4) inflammasome (FlaTox (15, 16)) and used it in primary murine IEC, instead of classically immortalized cell line experiments, to better approximate a physiologically relevant model. We developed an unbiased approach named Caspase Activity Profiling Through Unbiased Residue Evaluation, “CAPTURE”, to identify proteins cleaved by caspases. We analyzed the presence of aspartic acid (D) in peptides predicted to result from sequential cleavage, first by caspase, then by trypsin, compared to peptides observed in trypsin-only digestions. To further refine our analysis, we utilized a dual data reduction strategy, thereby reducing false positives by 90%. Applying this method in our experimental setup, we identified both previously characterized and uncharacterized caspase substrates, with several structural and junction proteins among the top hits. Selected targets were validated by Western blot. Finally, we demonstrate that our workflow can be applied to existing proteomics datasets, even those originally generated for unrelated experimental purposes, to examine potential caspase activity.

## Experimental Procedures

### 1. Primary epithelial cell culture and inflammasome activation

Crypts were isolated from the small intestine of mice and monolayer cultures were generated as previously described (17). Briefly, approximately 12 cm of proximal small intestine was isolated, and the villi scraped off with a glass slide. Following three PBS washes, the tissue was cut into 2mm sections, then incubated in 2mM EDTA for 30 minutes at 4 °C. After removing the EDTA solution, PBS was added, and the intestinal pieces were pipetted up and down with a 10mL serological pipet 3 times and then filtered through a 70μm strainer. 48 well plates were coated with 125 μL of a 2% Matrigel solution in cold DMEM and incubated at 37 °C for at least 30 minutes. Pelleted crypts were resuspended in complete organoid media (DMEM/F12 with Penicillin and Streptomycin, 1mM HEPES, 1x N2 and B27 supplement, 5mM N-acetylcysteine, 50mg/ml EGF, 1/20 Volume Noggin and R-spondin conditioned media) (18, 19) with 10 μM Y27632. Crypts were plated and incubated at 37 °C overnight. Media was replaced with complete organoid media without Y27632 and incubated for at least one hour prior to treatment. Recombinant proteins, protective antigen (PA) and N-terminus of lethal factor (LFn) fused to *Legionella pneumophila* flagellin (LFn-FlaA), were purified from *E. coli* as described before (16, 20). Endotoxin was removed using Pierce endotoxin removal columns. NAIP–NLRC4 Inflammasome activation was achieved by combining 4 μg/mL PA and 1 μg/mL LFn-Fla in media for 30 minutes at 37 °C. Cells were trypsinized for 5 minutes at 37 °C. Media was added to dilute out trypsin and the cells were centrifuged at 400xg for 5 minutes at 4 °C. Cells were washed with PBS twice and centrifuged at 400xg for 5 minutes at 4 °C then flash frozen in liquid nitrogen and stored at -80 °C until further analysis.

All mouse experiments were performed as approved by the Institutional Animal Care and Use Committee of the OHSU.

### 2. Organoid culture and Western Blot

Organoids were isolated and grown according to established methods (21). Organoids were grown in Matrigel for 1-2 days prior to exposure to FlaTox (16 μg/mL PA and 4 μg/mL Lfn-Fla) for 45 minutes. Samples were harvested from Matrigel domes using ice-cold Cultrex Organoid Harvesting Solution (R&D Cat No. 3700-100-01). 200μL was added to each well and up to 6 wells of confluent organoids were combined into an Eppendorf tube and kept at 4 °C with gentle rocking for 30 minutes, then centrifuged at 500xg for 5 minutes at 4 °C. Supernatant was removed and cells were lysed in RIPA Buffer (ThermoFisher), supplemented with protease inhibitor cocktail (Sigma). Lysates were vortexed, pelleted, and then kept at -20 °C. The following day, lysates were thawed and then spun at 10,000 xg for 10 minutes at 4 °C. Protein concentration was assessed by BCA. 60μg (25 μg for Cdh17) of sample was boiled for 10 minutes at 95 °C in sample buffer with DTT, loaded onto 4-20% Mini-PROTEAN TGX Precast Protein Gels (BioRad) and run at 180 V for 1 hour. The Trans-Blot Turbo Transfer System (Bio-Rad) was used to transfer to a 0.45 μm pore size PVDF membrane (Bio-Rad). The PVDF membrane was blocked for 1 hour at room temperature using Intercept (TBS) Blocking Buffer (927-60001, LI-COR). Primary antibody was incubated at 4 °C overnight in Intercept (TBS) Blocking Buffer with 0.01% Tween20. Primary antibodies and dilutions include: 1:500 β-actin (Santa Cruz Biotechnology; sc-47778), 1:1000 CASP1 (Cell Signaling; 24232), 1:1000 CASP7 (Cell Signaling; 8438), 1:1000 CASP8 (Cell Signaling; 8592), 1:10,000 CDH17 (RND Systems; MAB8524-SP), 1:1000 KRT20 (Cell Signaling; 13063T). The membrane was washed 3 times with TBS with 0.01% Tween20. Secondary antibody was incubated at room temperature for 1 hour in TBS with 1:10,000 Goat Anti-Rabbit HRP-conjugated (Sigma) or 1:10,000 Goat Anti-Mouse HRP conjugated (Sigma) and 1:2000 Rhodamine Anti-GAPDH Primary Antibody (Bio-Rad; 12004168). After 3 TBS + 0.01% Tween20 washes, proteins were detected by chemiluminescence with Clarity Max ECL substrate and imaged on a ChemiDoc (Bio-Rad).

### 3. Mass Spectrometry analysis

#### Sample preparation

Cell pellets from untreated and 30 minutes FlaTox treated cells were digested using a micro S-Trap™ micro spin column system (S-Trap^™^ Micro Column, Protifi) according to the manufacturer’s directions. Briefly, pellets were resuspended in 23 µL of 5% SDS + 50 mM triethylammonium bicarbonate buffer (TEAB) pH 8.5, vortexed, and sonicated for 5 minutes (in a bath sonicator). Then, proteins were reduced in 5 mM TCEP for 15 minutes at 55 °C at 800 r.p.m. and alkylated for 15 minutes at room temperature in the dark with 20 mM iodoacetamide. After protein alkylation, phosphoric acid was added to 2.5% final to pH<1, and 165 µL binding buffer of 100 mM TEAB pH 7.55 in 90% methanol was added and samples were added to the S-trap columns. Proteins were washed three times with binding buffer and digested overnight at 37 °C directly in the columns by adding trypsin (Promega) 1:10 (µg:µg) in a 20 µl final volume. Peptides were recovered by centrifuging the columns, adding 40 µL of 50 mM TEAB in water pH 8.5, 0.2% formic acid, and 50% acetonitrile in water consecutively. Peptides were dried down in a speed vac until fully dry and resuspended in 100 µL of trifluoroacetic acid (TFA) 0.1%, vortexed for 1 minutes and sonicated for 5 minutes. Samples were then applied to micro spin SPE columns prewashed by two volumes of 100% methanol and two volumes of 0.1% TFA. Samples were washed with four volumes of 95:5 TFA 0.1%: acetonitrile (ACN) and eluted with one volume of 80:20 ACN:TFA 0.1%. Peptides were dried down in a speed vac until fully dry and resuspended in 30 µL of MiliQ water for mass spec analysis.

#### Mass spec analysis

In all cases, peptides were fully dried in a speed-vac and resuspended in miliQ ultrapure water at 0.1 mg/mL (quantified by micro-BCA, Pierce) before LC-MS/MS analysis. Peptides were analyzed by 5 µL injection volume on a Waters NanoAquity UPLC system (Waters Corporation, Milford, MA, USA) with a custom-packed C18 column (20cm x 25 μm inner diameter Phenomenex Jupiter) coupled with a Q-Exactive mass spectrometer (Thermo Scientific, San Jose, CA, USA). Peptide separation was carried out in a C18 reverse-phase analytical column with the following solvents (A) 0.1% formic acid (B) ACN with 0.1% formic acid over 120 minutes. Eluting peptides were directly analyzed by nanoelectrospray ionization and full-MS full scans were collected at scan range 300–1800 *m/z* at a resolution of 70,000 in positive mode. The top 12 most intense parent ions were submitted to high-energy collision-induced dissociation (HCD) fragmentation (1.5 *m/z* isolation width; 30% normalized collision energy; 17,500 resolution at 400 *m/z*), before being dynamically excluded for 30 s.

#### Data identification and statistical analysis

Raw spectra were transformed into mzML using MSConvertGUI (Version: 3.0.24239-76e520 and e) and analyzed in FragPipe (22) v22.0 using data dependent configurations by matching against the mouse reference proteome database from the Uniprot database UP000000589. Fragpipe search engine “Protein digestion” parameters were set as follows: semi-specific tryptic cleavage (e.g., after K or R) was authorized with a maximum of 3 miscleavages on the C-terminus side of the peptides, caspase (e.g., after D) was authorized with a maximum of 5 cleavages on the C-terminus side of the peptides. The ion mass tolerance was set at 20 PPM for both precursor and fragment ions. The data were filtered at 1% false-discovery rate in both peptide-spectrum matches and protein levels. Protein quantification was performed by data-dependent analysis (DDA) using label-free quantification (LFQ) with default parameters. The resulting data was further processed using the R package Romics Processorv1.1 (https://github.com/PNNL-Comp-MassSpec/RomicsProcessor).

The data were log2-transformed and median-normalized, then statistically analyzed by Student’s T-test to compare the protein abundance between conditions and G-binomial test to compare presence/absence of peptides in the different groups. In all cases, only proteins or peptides present in at least 50% of replicates of one group were considered for analysis.

Imputation was only used for peptide heatmap visualization and KEGG pathway enrichment analysis in **Supplementary Figure 1**.

## Results

### Short-term inflammasome activation has a negligible effect on protein abundance

To analyze CASP cleaved targets in primary epithelial cells, we took advantage of recently developed protocols to isolate and culture primary epithelial cells from the mouse intestine (17, 23). We isolated crypts from the murine small intestine and cultured them as monolayers to treat with FlaTox to activate the NAIP-NLRC4 inflammasome. FlaTox is a well-characterized tool for the delivery of flagellin directly into the cytosol, which leads to rapid and specific induction of NAIP-NLRC4 inflammasome assembly, CASP1 activation, and pyroptosis (9, 16). Cells that were treated with FlaTox for 30 minutes and untreated controls were flash frozen. Thus, we generated protein samples from primary cells with and without inflammatory caspase activation (**Fig. 1A**).

**Figure 1.**
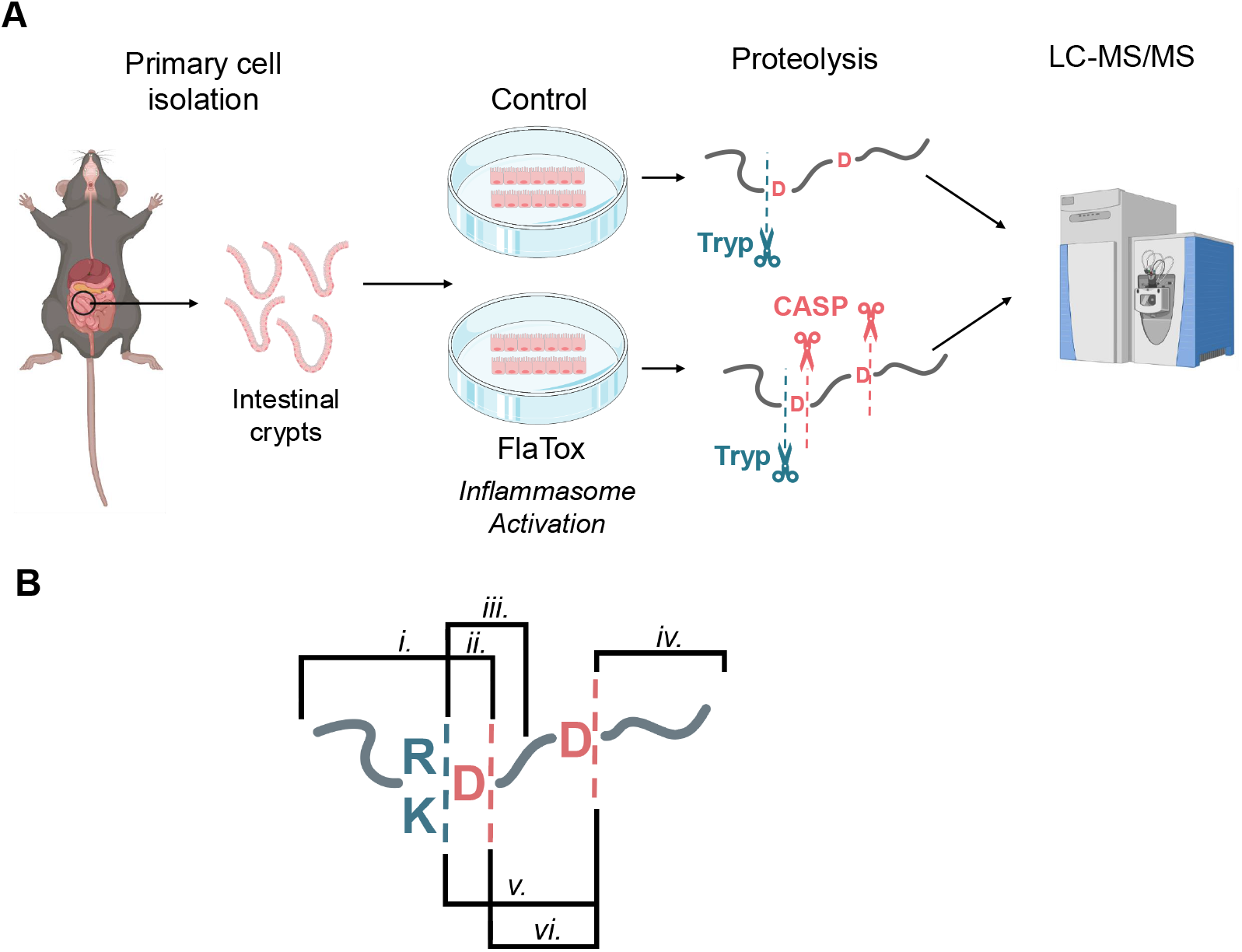
Experimental design. **A**. Primary epithelial cells were collected from murine small intestine and cultured under control conditions or treated with FlaTox for 30 minutes to induce inflammasome activation. Following treatment, cells were lysed and proteins digested with trypsin (Tryp). Peptides were analyzed by LC-MS/MS and identified using Fragpipe v22.0. **B**. Representation of all possible CASP-derived peptides from a given protein, sequence required to be considered a potential CASP-derived peptide are indicated as **i**. Arginine (R) or Lysine (K) at C-ter, followed by aspartic (D); **ii**. D at N-ter; **iii**. D before C-ter (for practical purposes, equivalent to **v**.) **iv**. D at N-ter and C-ter are also preceded by R or K.

Proteome analysis from whole cellular extracts resulted in a total of 6,329 proteins identified (**Supplemental Table 1**). From those, five proteins: IgGFc-binding protein (Fcgbp), calcium-activated chloride channel regulator 1 (Clca1), zymogen granule membrane protein 16 (Zg16), mucin-2 (Muc2) and glutathione S-transferase A5 (Gsta5) were decreased in abundance in the FlaTox treated cells and no proteins were increased in FlaTox treated cells, indicating a minor effect of the treatment on protein abundance (**Supplemental Fig. 1A, Supplemental Table 1**). Since FlaTox treatment had a negligible impact on protein abundances, we focused on the peptide levels to determine potential alterations as a consequence of inflammasome activation. From a total of 57,558 peptides identified (**Supplemental Table 1**), 1,394 were significantly more abundant in the FlaTox treated cells and 1,987 were significantly more abundant in the control cells (**Supplemental Fig. 1B, Supplemental Table 1)**. We expect that peptides that are less abundant following inflammasome activation represent caspase targets before cleavage.

Next, we analyzed their precursor proteins by KEGG pathway enrichment analysis to evaluate the overall effect of inflammasome activation. From a total of 83 pathways enriched in control samples and 60 enriched with inflammasome activation, 39 were found in common (**Supplemental Table 1**). Both unique and shared KEGG-enriched pathways include entries for general metabolism, **(Supplemental Fig. 1C**). Overall, the analysis of proteins or peptides did not reveal a clear signature of caspase activity, or an induction of pathways involved, indicating that the effects of inflammasome activation or caspase activity cannot be detected using a tryptic LFQ workflow alone.

### CAPTURE strategy reveals potential caspase-target peptides overlooked by conventional LFQ analysis

While previous studies have mainly used N-termini labeling methods to identify caspase targets (6, 7) requiring relatively complex sample preparation, here we investigated whether a standard bottom-up LFQ data-dependent analysis workflow, combined with refined data analysis, could detect potential caspase targets based on the peptide sequences.

Upon a close inspection of known caspase targets, we could not identify a clear motif sequence for caspase cleavage sites. Instead, the only common factor among caspases 1-10 is that the cleavage site is at the C-terminal end of the peptide, following an aspartic residue (D) (**Fig 2A)**. Therefore, we designed the Caspase Activity Profiling Through Unbiased Residue Evaluation (CAPTURE) strategy. This approach leverages the presence of D at the C-termini of the cleavage site to detect the presence of potential caspase-derived peptides. As a result, five theoretical peptide types (see **Fig. 1B**) can be generated by the combined cleavage specificities of caspases and trypsin, the most used enzyme for bottom-up LFQ pipelines across mass spectrometry proteomics laboratories. Considering all five theoretical peptide types, we detected a total of 5,686 potentially caspase-derived peptides (**Supplemental Table 1)**. Continuing with the selection of potential CASP-target peptides, we applied a dual data-reduction strategy. First, we discarded all peptides originating from proteins exhibiting significant abundance changes upon treatment (Fcgbp, Clca1, Zg16, Muc2, Gsta5). Next, we performed a two-step statistical assessment evaluating peptide-level abundance differences between conditions using a Welch’s T-test, and presence/absence patterns of peptides across conditions using a G-binomial test. Our data-filtering workflow resulted in a final list of 432 CASP-derived peptides candidates, reducing the initial set of potential CASP-derived peptides by ∼92% of likely false positives. From those, 223 were increased in the control and 209 increased after inflammasome activation (**Fig. 2B, Supplemental Table 1**). To explore if quantitative analysis was representative of any biological meaning, we examined their parental proteins using KEGG pathway enrichment. A total of 19 pathways were enriched in the control condition and 22 were enriched upon inflammasome activation, again showing comparable numbers between conditions but with notable qualitative differences. In this case only 6 enriched pathways were found in common, mainly related to general metabolism (**Supplemental Table 1**). We detected that FlaTox treatment led to an increase in potential caspase-derived peptides, predominantly associated with amino acids and protein processing, nucleotides, and cell adhesion, suggesting that these processes may be disrupted. Instead, controls also contained potential caspase-derived peptides, primarily from proteins involved in carbon metabolism and redox/detoxification processes, likely reflecting basal caspase activity (**Fig. 2C, Supplemental Table 1**).

**Figure 2.**
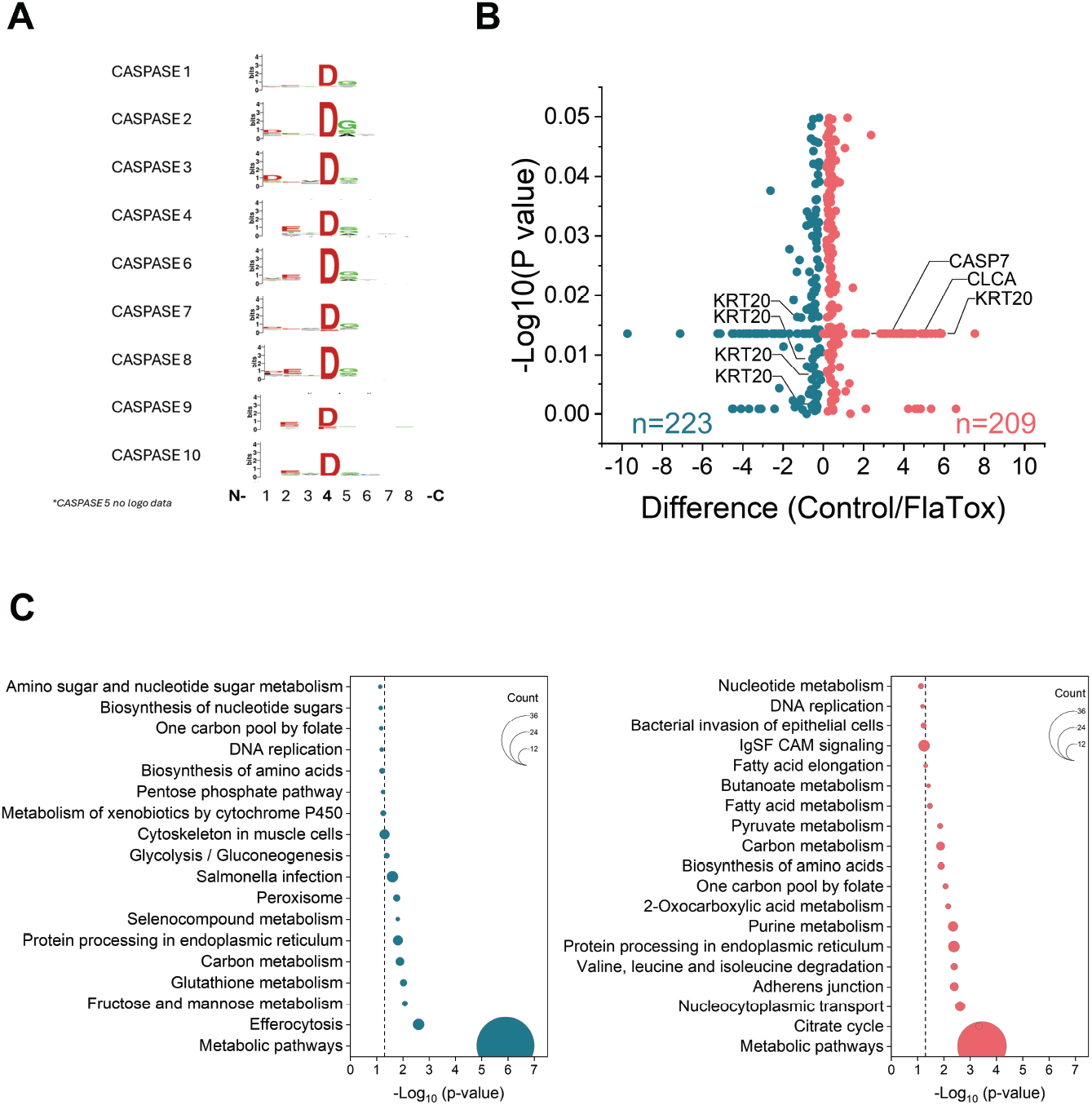
Selection of significant peptides with the potential of being CASP-derived. **A**. Logo sequence from caspase 1-10. Negative amino acids (in red), basic amino acids (in green) and neutral (in black). **B**. Volcano plot of significant CASP-likely peptides due to FlaTox treatment (in red) and control (blue). Parental proteins from significant peptides were analyzed in DAVID and pathway enrichment is shown in **C**.

### Known and novel Caspase targets were detected using the CAPTURE strategy

To determine whether the potential caspase-derived peptides originated from bona fide caspase substrates, we first compared our list with previously reported caspase targets. Notably, CASP7, a well-described substrate of CASP1, was present among FlaTox-induced caspase-derived peptides(7). Specifically, the CASP-derived peptide 197(D)SGPINDIDANPR210 was only detected upon inflammasome activation (*p* = 0.018). Importantly, this peptide cannot result from trypsin cleavage alone; instead, trypsin digestion of CASP7 can generate the longer peptide 186(R)GTELDDGIOADSGPINDIDANPR210, which contains the 197–210 sequence. The longer peptide was increased in control samples (*p* = 0.016) (**Fig. 3**). Together, these findings suggest that in control samples, CASP7 was cleaved only by trypsin, whereas in FlaTox-treated samples, a caspase-generated precursor peptide exists prior to tryptic digestion. We confirmed this observation by Western blot in an independent experiment, detecting a unique band at 20 kDa after FlaTox treatment of primary epithelial organoids **(Fig. 4A**). Also, Caspase-7 was recently shown to be required for the final stages of extrusion of intestinal epithelial cells (13), indicating that our dataset contains proteins that are crucial for the extrusion process.

**Figure 3.**
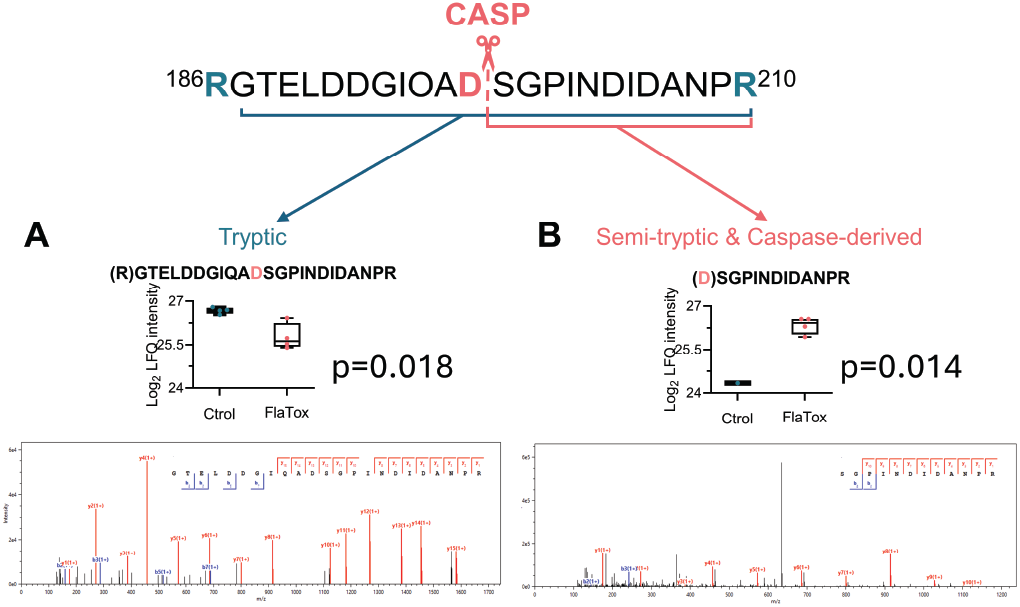
Caspase-7 peptides detected by CAPTURE method. **A**. CASP7 tryptic peptide identified by the LFQ analysis. **B**. Potentially caspase derived peptide detected only in FlaTox treated cells, identified after CAPTURE method.

**Figure 4.**
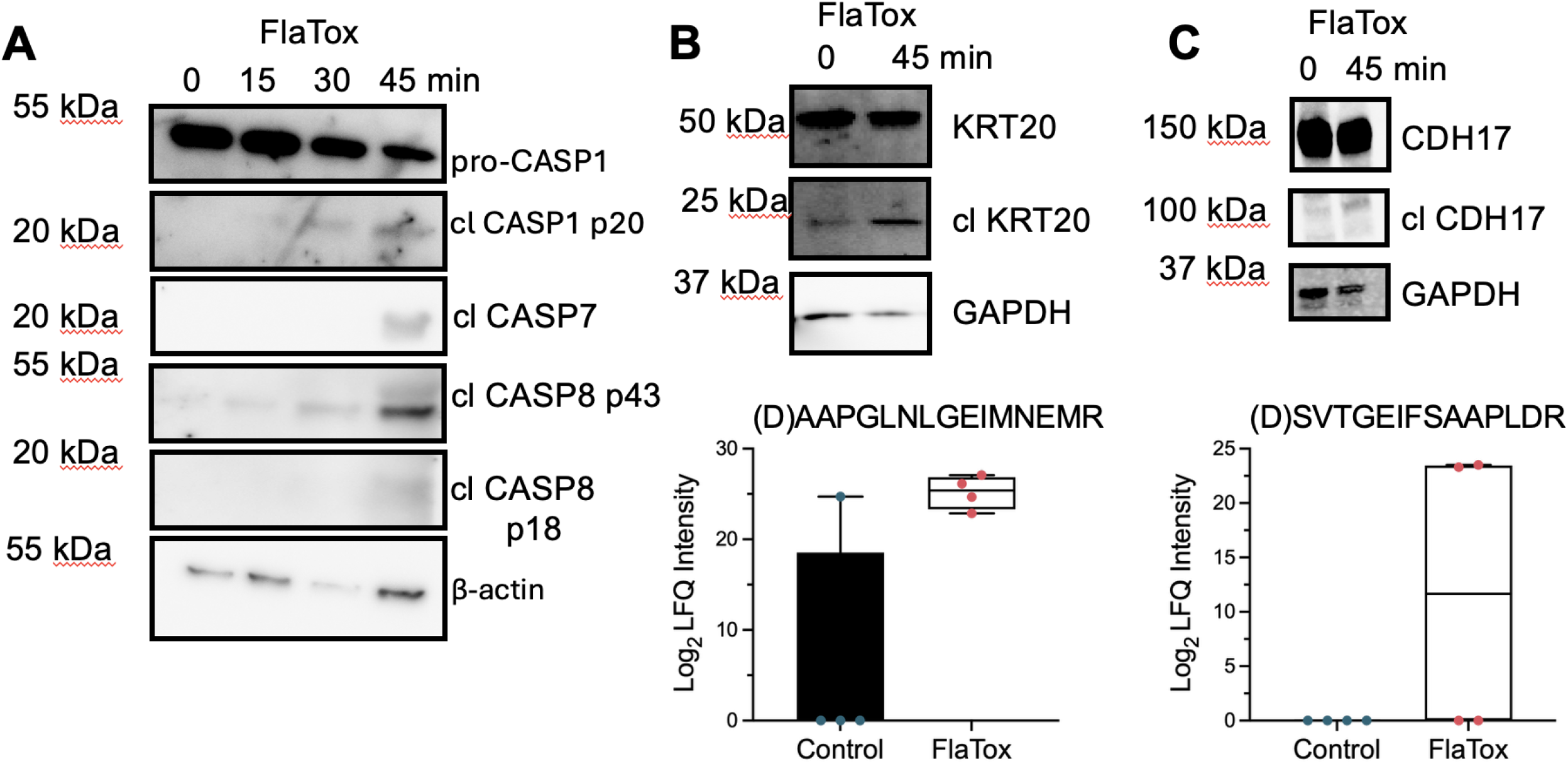
Potential novel CASP-targets validated by Western blot. **A**. Cleavage of CASP1, CASP7, and CASP8 following inflammasome activation. **B**. Organoids treated with FlaTox for 45 minutes show an increase in the presence of an approximately 25kDa band representing the cleavage of keratin 20 (KRT20). **C**. Organoids treated with FlaTox for 45 minutes show an increase in the presence of an approximately 100kDa band representing the cleavage of cadherin 17 (CDH17).

In addition, we confirmed by Western blot the cleavage of Keratin20 (KRT20) (**Fig. 4B**). While cleavage of KRT20 was previously shown under apoptosis-inducing conditions (24), we show here that this cleavage occurs in response to inflammasome activation as well. No previous studies of caspase targets have identified KRT20, most likely because none were conducted in intestinal epithelial cells.

Once we were able to identify known caspase targets using CAPTURE, we next aimed to define previously unknown caspase targets. Cadherins are calcium-dependent adhesion molecules that are highly expressed in the intestinal epithelium and play a critical role in maintaining barrier integrity (25, 26). E-cadherin, known to be cleaved by CASP3 (27), was one of our top hits, but we also found several potential CASP-cleavage peptides from Cadherin17, where the enrichment of peptides did not reach significance (*p*=0.06), possibly due to the variability between samples. Western blot confirmed that Cadherin 17 (CDH17) was cleaved, generating an approximately 100kDa fragment (**Fig. 4C**). Overall, these results indicate that our method enables the identification of new caspase targets in primary cells and that inflammasome activation in IEC leads to cleavage of many structural proteins.

### The CAPTURE strategy can be used for Caspase activity detection by reanalyzing LFQ datasets

To investigate whether the CAPTURE strategy could be retrospectively applied to previously generated LFQ datasets obtained using a different approach we applied our method to a public dataset. We used the repository PXD021501(28) where MIN6 cells were exposed to pro-inflammatory cytokines and corresponding controls, since TNFα treatment can activate apoptotic caspases via the extrinsic pathway (29, 30). From a total of 48,439 peptides identified, 4,138 represented potential CASP-derived peptides based on their sequences (**Supplemental Table 2)**. After applying the CAPTURE strategy 512 peptides were increased in cytokine treatment while 1,338 represented differential levels in the controls (**Supplemental Table 2)**. When we analyzed the biological relevance of those peptides by KEGG pathway enrichment found 50 pathways enriched under cytokine treatment, and 48 pathways in controls (**Supplemental Table 2)**. Importantly, many of the cytokine-associated pathways were consistent with known or suspected caspase activity, including proteasome function, mitochondrial metabolism, and canonical apoptosis pathways, whereas enrichment in controls was largely limited to nonspecific cellular responses (**Fig. 5**). By applying CAPTURE, we detected 31 peptidyl sequences matching between the list of CASP-likely peptides from FlaTox treatment (current study) and those also resulting as CASP-likely peptides as consequence of cytokine treated cells, providing strong evidence as peptide derived from unbiased caspase action. Additionally, among them are known caspase targets including, filamin B (31), myosin-9 (32), nuclear mitotic apparatus protein 1 (33), and serine/arginine rich splicing factor 1 (34).

**Figure 5.**
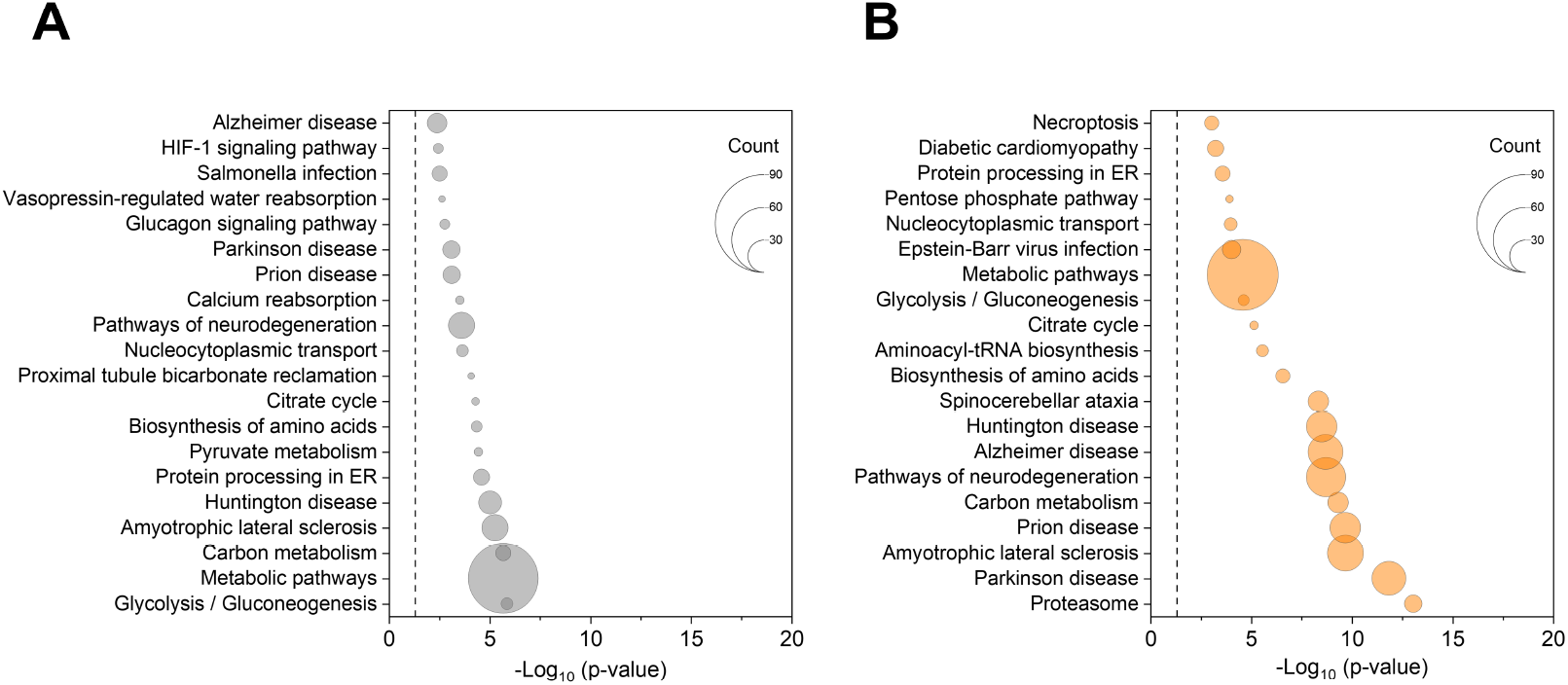
Bubble plot of KEGG pathway enrichment of CASP-likely peptides from cytokine treatment in MIN6 cells. CAPTURE was applied in a public dataset and peptides increased in cytokine treatment were evaluated by KEGG pathway enrichment analysis. Top 20 pathways enriched for controls (Gray) and cytokine (Orange) treatments (**Supplemental table 2**).

## Discussion

Detection of caspase targets has traditionally relied on approaches such as western blot or mass spectrometry-based workflows that require enrichment or labeling steps. While these methods have successfully identified new caspase targets, they have inherent limitations, including the need for expensive reagents and sophisticated workflows, which may be restrictive for non-specialized laboratories. Moreover, the number of targets identified using these approaches, although significant, remains limited. In this study, we hypothesized that a standard label-free quantification (LFQ) workflow could facilitate the detection of multiple caspase targets without requiring additional reactions or labeling steps. FlaTox is a potent inducer of caspase-1 activation (15, 16). Taking advantage of this system, we analyzed full extracts from mouse intestinal epithelial cells to identify potential caspase-derived targets. Here, we demonstrate for the first time that a standard DDA LFQ approach can be effectively used to detect caspase substrates with high confidence, reducing the number of potential false positives by ∼90%. Additionally, we applied our method to an independent dataset, detecting potential enrichment of caspase-derived peptides, thus extending the applicability of this approach to other experiments, even those not designed to address CASP activity. Using primary intestinal epithelial cells, we observed activation of CASP1, which led to the cleavage of CASP7, a known CASP1 target. Notably, the peptide (<u>D</u>)SGPINDIDANPR was detected only upon inflammasome activation whereas the corresponding precursor peptide (R)RGTELDDGIOA<u>D</u>SGPINDIDANPR was increased in control conditions, indicating cleavage of CASP7 occurs specifically upon FlaTox-induced inflammasome activation. These findings were confirmed both at the MS2 spectral level and by Western blot, providing strong and robust validation of our observations (**Fig. 3 and 4A**). We also identify cleavage of KRT20 (**Fig. 4B**), which had been previously shown as cleaved upon induction of apoptosis, but we identify that this cleavage also occurs with pyroptotic stimuli. Likely, KRT20 was not found in previous studies of pyroptotic caspase targets, as none have been conducted in epithelial cells.

Three previous studies have examined CASP1 targets directly or used pyroptotic stimuli to examine caspase substrates; all of them were conducted in monocytes and macrophages, with none utilizing primary cells. Our data shows limited overlap with previous studies: there are 10 proteins in common with Shao et al. 2007, 5 proteins in common with Lamkanfi et al. 2008, and 7 proteins in common with Agard et al. 2010. While cell type would appear to be a likely driver, the fact that our CAPTURE method found 432 potential caspase substrates while previous studies found fewer than 60 is the most likely cause for the lack of overlap between datasets. Inflammasome activation in intestinal epithelial cells canonically leads to Gasdermin D cleavage and cleavage of IL-18. Here, we identify cleavage of additional caspase substrates.

Limitations of the study

1. NAIP–NLRC4 activation leads primarily to CASP1 activation. CASP8 can also be recruited and activated via the adaptor ASC, but this is more pronounced in the absence of CASP1 (35). Further, CASP8 deficiency only has an effect in CASP1 deficient animals in a model of oral *Salmonella* infection (9, 36), suggesting CASP1 is the dominant Caspase activated. Finally, we find CASP7 robustly cleaved in our samples, which also cleaves many downstream targets. We tried to limit this caveat by choosing a timepoint of 30 minutes where we observe primarily CASP1 cleavage (**Fig. 4A**).
2. We found a similar number of potential CASP-derived peptides in control and in FlaTox treatment. While some control peptides may indicate basal caspase activity, some are a consequence of the CAPTURE method. For example, two peptides of CASP1 are increased in the control. The peptide 300(D)SEEDFLTDAIFED**D**GIKK includes the cleavage site at D316 which releases the p10 subunit of CASP1 (37, 38), treatment with FlaTox and subsequent caspase activation would lead to a decrease in this peptide. The second peptide 324(D)FIAFCSSTPDNVSWR is part of the p10 fragment, which can be released from the cell through the pores due to the lytic nature of pyroptosis. Sampling the supernatant as well as whole cell lysates could further inform this analysis. Additionally, experiments could be conducted in GSDMD deficient cells to prevent loss of peptides through the pores.
3. Neither GSDMD nor IL-18 were among the CASP-derived peptides we found using CAPTURE. No peptides that include the known cleavage sites of these proteins were detected. Previous studies on CASP1 targets in myeloid cell lines used neo-N-terminus labeling to identify proteins, though no previous study has ever managed to identify CASP1, GSDMD, and IL-18 (IL-1β in myeloid cells) all at once. Development of pre-enrichment steps may be a powerful complement in the future.
4. Even after a reduction of more than 90% using various filtering strategies to limit false discovery, some of the remaining candidates may still be false positives. The post-aspartic acid cleavage motif that we use in this study could also be the result of additional non-caspase-like proteases. Therefore, we recommend the use of our approach as a method to create a preliminary dataset that still requires further exploration and/or cross-validation with other methods. For instance, we found several proteins that are already known to be cleaved by caspases such as E-Cadherin (27), Vimentin (39), KRT18 and KRT19 (40), or Plectin (41). Additionally, there are some potential CASP targets that did not reach the level of significance, such as CDH17. Increasing the number of replicates could improve statistical power and provide a more robust dataset for further validation.

Despite these limitations, we identified new cleaved proteins post-inflammasome activation, notably several cytokeratins and other structural proteins. Cytoskeleton rearrangement is a requirement to drive rapid cell extrusion of dying IEC (11, 12). The cleavage of cytokeratins is likely required for the cell to change shape, while the cleavage of cadherins may be necessary to allow the cell to detach from its neighbors and be removed from the tightly packed epithelium. Functional assays mutating the cleavage sites in these proteins would be needed to demonstrate the role in extrusion. This dataset can be the basis for the design of such future experiments. Additional cell types, such as keratinocytes and endothelial cells, warrant further study to examine how inflammasome activation leads to cleavage of a specialized subset of targets in those cells. This could lead to a better understanding of the unique outcomes of pathogen sensing in different organs and tissues.

## Abbreviations

CAPTURE: (Caspase Activity Profiling Through Unbiased Residue Evaluation)
CASP: (Caspase)
DDA: (data-dependent analysis)
IEC: (intestinal epithelial cell)
LFQ: (label-free quantification)

## Data Availability

The original mass spectra have been deposited on the public proteomics repository MassIVE and are accessible at MSV000098956.

## Acknowledgements

Partially supported by the PMedIC project through the Laboratory Directed Research and Development Program at Pacific Northwest National Laboratory, a multiprogram national laboratory operated by Battelle for the US Department of Energy under contract number DE-AC02-05CH11231. This work was supported by the National Institutes of Health, National Institute of Allergy and Infectious Diseases (R01 AI167974) and a Collins Medical Trust award to I. Rauch. A.R. Gibson is supported by the Howard Hughes Medical Institute Hanna H. Gray Postdoctoral Fellowship.

## Author Contributions

A.R. Gibson: Conceptualization, Investigation, Writing-original draft, Funding acquisition I. Diaz Ludovico: Investigation, Methodology, Formal analysis, Visualization, Writing-original draft

G. C. Clair: Methodology, Writing-review and editing

C. M. Hutchinson-Bunch: Investigation

J. Adkins: Writing-review and editing, Funding acquisition

I. Rauch: Conceptualization, Writing-review and editing, Funding acquisition

## Supplementary Data

**Supplemental Figure 1. Proteomic analysis after FlaTox treatment**.

**A**. Proteins identified in Fragpipe were statistically analyzed and represented in a volcano plot. Increased proteins in control are indicated in blue. **B**. Heatmap of total analyzed peptides indicating those with significant changes as consequence of FlaTox treatment. **C**. Top 15 pathways of parental proteins from peptides indicated in **B**.

**Supplemental Table 1**

FlaTox peptidomics and KEGG analysis

**Supplemental Table 2**

MIN6 peptidomics and KEGG analysis

