## Supplementary figures and images for "Unbiased proteomics following inflammasome activation identifies caspase targets in primary intestinal epithelial cells"

### Supplemental figure 1

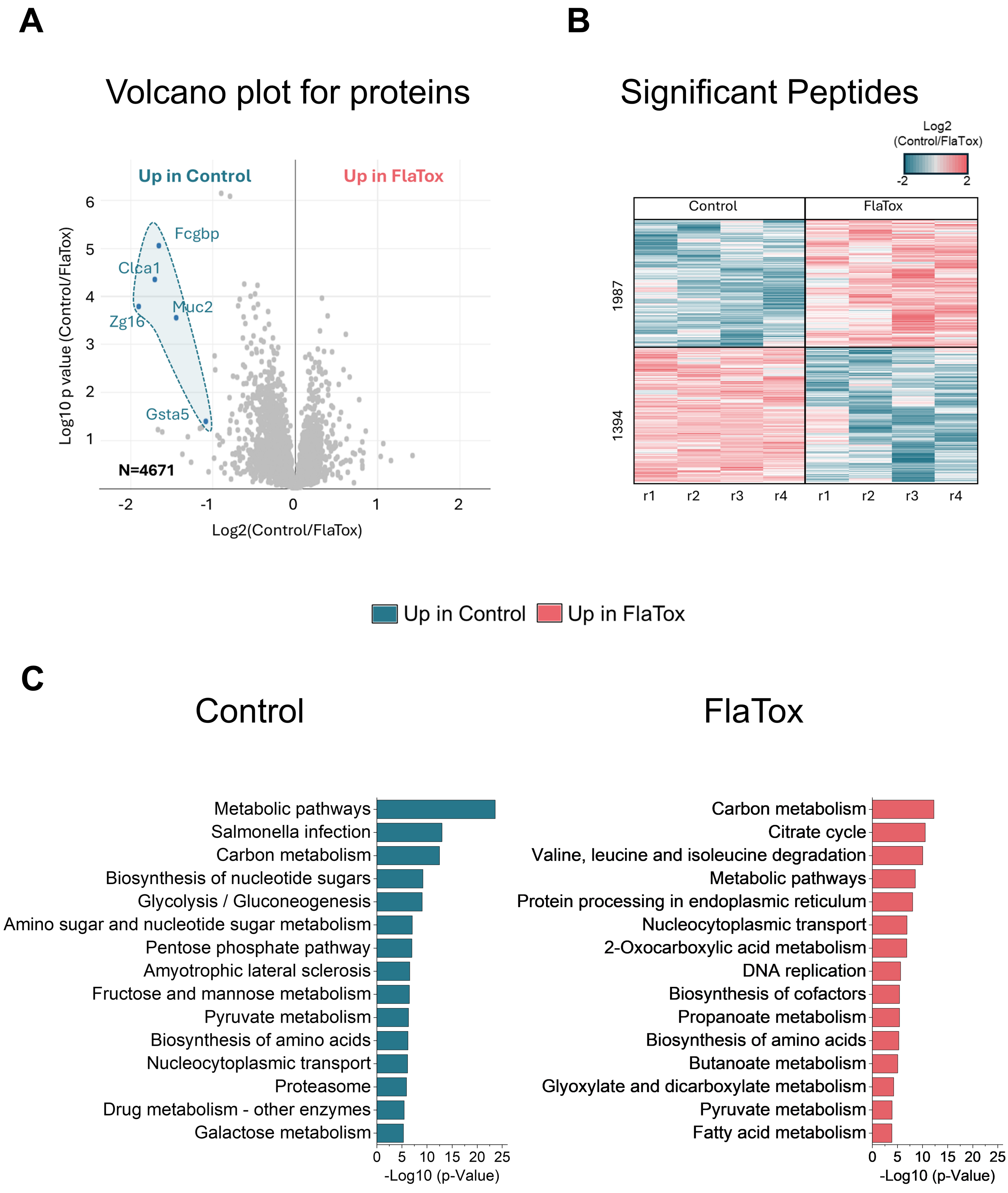
